# Graph-theoretic comparisons of structural covariance networks: quantifying the false discovery rate

**DOI:** 10.64898/2026.06.23.733999

**Authors:** James Read-Tannock, Andrew T. Reid, Etienne Farcot, Martin Schürmann, Christopher R. Madan

## Abstract

Structural covariance networks (SCNs) represent spatial patterns of covariation in brain morphology, often as a network of connections between nodes representing correlations in grey matter volume or cortical thickness measured by magnetic resonance imaging (MRI). SCNs have been suggested to reveal differences in functional organisation that are reflected in coordinated alterations to brain structure, and often these differences are sought in graph-theoretic measures such as the degree of clustering, segregation into distinct modules, or the characteristic path length between nodes. A common practice is to calculate SCNs for groups of interest, and use permutation testing to determine if they are significantly different for the measure of interest. However, the statistical validity of group comparisons using SCN-derived graph measures remains poorly understood.

Here, we systematically evaluate the reliability of SCN estimation and downstream graph-theoretic analyses using structural MRI data from the Human Connectome Project (*n* = 1,096). We use simulations to show the effects of sample size and atlas dimensionality on SCN reliability. Using bootstrapping to characterise the distribution of SCN graph measures, we establish that small sample sizes systematically bias graph-theoretic measures including clustering, characteristic path length and modularity. Finally, we use simulations based on extrema from the bootstrapping distribution to characterise the statistical power and false discovery rate (FDR) for graph-theoretic between-group comparisons of SCNs, showing that at small sample sizes (*n* ≤ 30) permutation testing is no better than chance.

These findings suggest that many significant SCN group differences, particularly those using small samples and high-dimensional parcellations, may reflect sampling noise rather than true biological differences. We recommend that future SCN studies use larger samples, coarser parcellations, and explicitly evaluate reliability before interpreting group differences.

## Introduction

The human brain has long been understood as a complex network of interacting, functionally specialised structures (Mesulam, 1998), and in recent years this connectivity structure has begun to be characterised using the mathematical framework of graph theory. In this framework, brain regions are represented as distinct ‘nodes’ and the relationships between them as ‘edges’, allowing the macro-scale connectivity structure observed with magnetic resonance imaging (MRI) to be quantified in terms of graph-theoretic measures (henceforth ‘graph measures’) such as clustering, small-worldness, and node centrality (Bassett & Sporns, 2017; Bullmore & Sporns, 2009). These measures describe how the pattern of connections in a network determine its information passing properties, and thus how the integration and segregation of specialised brain regions may subserve various cognitive demands (Cohen & D’Esposito, 2016; Sporns, 2013). For this reason, graph measures of brain networks have become an object of study in cognitive neuroscience and have been linked to individual differences in cognition, development, and disease.

Structural covariance networks (SCNs) apply these ideas to patterns of covariation in brain morphology. SCNs are constructed by correlating regional measures of grey matter morphology (e.g., cortical thickness or surface area) across individuals, revealing how brain anatomy may co-vary within a population (Evans, 2013; He et al., 2007). This approach requires only standard structural MRI data and thus provides an accessible route to large-scale network analysis. Graph-theoretic analyses of SCNs have reported associations with age, intelligence, and psychiatric and other clinical conditions such as schizophrenia, autism, and irritable bowel syndrome (Bethlehem et al., 2017; Hosseini et al., 2013; John et al., 2017; Khundrakpam et al., 2017; Labus et al., 2014; Palaniyappan et al., 2019; Singh et al., 2013; Wu et al., 2012). These findings have fuelled interest in SCNs as a means of identifying behaviour-associated variation in grey matter structure that has been hard to discover otherwise (Avinun et al., 2020; Kharabian Masouleh et al., 2019; Marek et al., 2022), and the use of SCNs has grown rapidly over the last two decades across many subfields, with group comparisons among the most common applications.

Commonly, differences in graph measures between groups are assessed for statistical significance by using permutation testing: exchanging observations between groups to produce a null distribution of the graph measure. However, despite a growing volume of publications, there has been limited investigation into the reliability and statistical validity of this method when applied to graph measures derived from SCNs. Studies often report significant effects using relatively small samples, but it remains unclear how often such differences could arise by chance. Previous work examined differences in correlation matrices between matched healthy control groups and found substantial variation depending on site of acquisition and processing pipeline, suggesting instability of SCN-derived graph measures (Carmon et al., 2020). Furthermore, community detection algorithms produced less stable solutions on SCNs compared to networks from other imaging modalities (Reid et al., 2016). Yet, no systematic evaluation of statistical power and false discovery rate (FDR) for graph-theoretic comparisons of SCNs across sample sizes has been conducted. This is a critical research gap: if permutation testing fails to control FDR at typical sample sizes, a substantial proportion of published SCN findings may represent false positives rather than true biological differences.

Here, we address this gap by estimating the statistical power and FDR obtained when testing for group differences in SCN graph measures, using the large Human Connectome Project (HCP) dataset. We use resampling methods to estimate population distributions for common graph measures, comparing results for large versus small samples, and conducting simulations to assess how sample size *n* and the number of atlas regions (atlas dimensionality) *p* influence error rates in downstream analyses. Our aim was to establish what sample size is needed for meaningful between-group differences in SCN graph measures. We predicted that, at low *n* and high *p*, permutation testing will have low statistical power and hence fail to control FDR in the published literature, since the number of pairwise correlations to be estimated grows quadratically with the number of regions. To our knowledge, this is the first systematic characterisation of statistical power and false discovery rate for graph-theoretic comparisons of SCNs across the range of sample sizes and parcellation resolutions used in practice.

Finally, it should be noted that many types of SCN have been developed. These include methods based on seed-based covariance or independent component analysis (typically not graph-theoretic; e.g. (Xu et al., 2009)), several methods for intra-individual SCNs based on the similarity of regional grey matter morphology (Kong et al., 2015; Tijms et al., 2012; Wee et al., 2012), individual differential SCN (IDSCN; (Liu et al., 2021)), morphometric similarity networks using multivariate correlations (Seidlitz et al., 2018), morphometric inverse divergence networks using multivariate Kullback-Leibler divergence (Sebenius et al., 2023), and causal SCNs using Granger causality on observations ordered by a variable of interest (e.g., disease severity; (Zhang et al., 2017)). In the present work, we only consider one prevalent approach to group SCN analysis first exemplified in (He et al., 2008) where the network is based on the sample correlation matrix of regional grey matter morphometric variables (such as cortical thickness or surface area), and differences in graph measures are assessed for significance using permutation testing.

## Methods

### Dataset

#### Cortical thickness

Structural MRI data were collected as part of the Human Connectome Project (HCP) at Washington University in St. Louis (Van Essen et al., 2012). Images were acquired on a Siemens Skyra 3T scanner equipped with a 32-channel head coil. High-resolution T1-weighted (T1w) images were collected using a 3D MPRAGE sequence (Mugler & Brookeman, 1990) with 0.7 mm isotropic resolution (FOV = 224 mm; matrix = 320; 256 sagittal slices; TR = 2400 ms; TE = 2.14 ms; TI = 1000 ms; flip angle = 8°; bandwidth = 210 Hz/pixel; GRAPPA = 2; 10% phase oversampling). T2-weighted (T2w) images were acquired with a variable flipangle turbo spin-echo (Siemens SPACE) sequence (Mugler et al., 2000) with identical spatial parameters (TR = 3200 ms; TE = 565 ms; bandwidth = 744 Hz/pixel; GRAPPA = 2). T1w/T2w volumes were acquired twice during the same session and averaged to improve signal-to-noise ratio. All images were subjected to the HCP quality control procedure (Marcus et al., 2013); only scans rated ‘good’ or ‘excellent’ (scores of 3 or 4 on a four-point scale) were retained. As a result, not all participants had both T1w and T2w volumes available for averaging.

Structural images underwent the HCP minimal preprocessing pipeline (Glasser et al., 2013), which uses FSL tools (Jenkinson et al., 2012) and the custom FreeSurfer (Fischl, 2012) v5.3-HCP protocol (https://github.com/Washington-University/HCPpipelines/blob/master/FreeSurfer/FreeSurferPipeline-v5.3.0-HCP.sh) to perform bias correction, distortion correction, and surface reconstruction using both T1w and T2w data.

We used the preprocessed structural data from the ‘HCP_S1200_GroupAvg_v1’ Dataset, comprising 1,096 participants (a subset of the 1,113 total participants with 3T structural MRI data). Specifically, we used cortical thickness structural data represented on the ‘fs_LR32k’ mesh (a left-right symmetric version of FreeSurfer’s ‘fsaverage’ template space) which has 32,492 vertices per hemisphere.

#### Cortical atlases

We analyzed a range of widely used cortical atlases in the form of surface parcellations spanning coarse to fine spatial scales. Hemispheres were treated separately (left/right parcels are distinct nodes), and subcortical structures were not included. For each atlas, we averaged vertex-wise cortical thickness within each parcel to obtain one value per parcel per subject. The set of atlases was chosen to cover approximately 10–300 nodes so that we could study how the atlas dimensionality impacts SCN reliability.

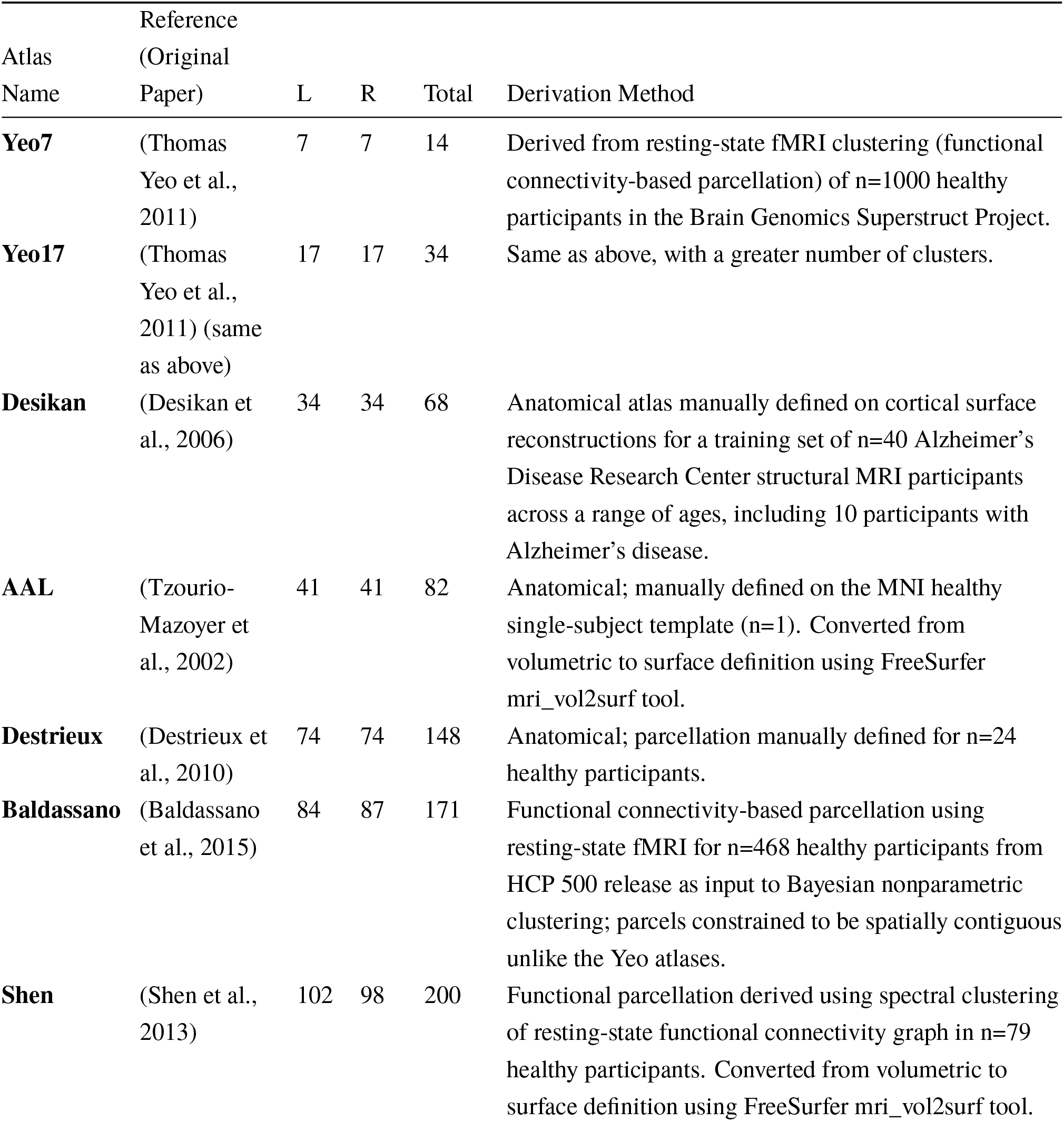
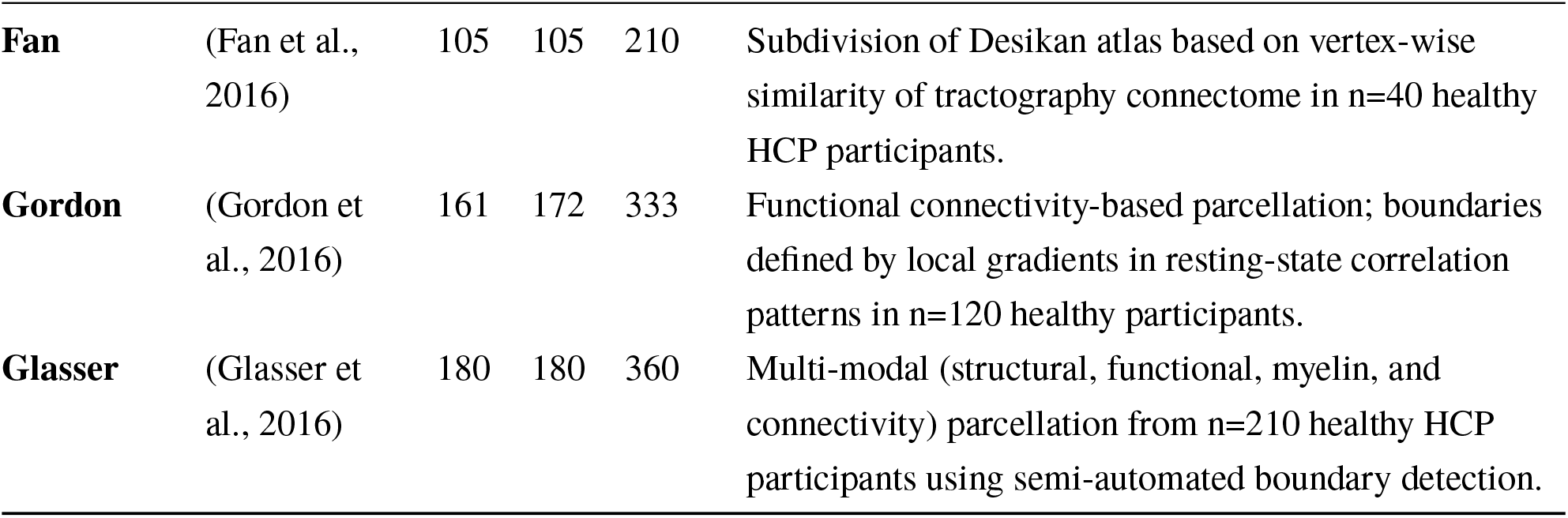

### Graph theory basic definitions

A graph *G* = (*V, E*) consists of a set of nodes *V* (also called vertices) and a set of edges *E* ⊆ *V* ×*V* representing the pairwise relationships between nodes. We denote the number of nodes |*V* | = *p* and the number of edges |*E*|. In the context of network neuroscience, nodes typically represent brain regions defined in an atlas (so *p* is the total number of brain regions or ‘atlas dimensionality’) while edges encode structural or functional connectivity between those regions. In SCNs, the ‘connectivity’ of two regions is defined using a measure of their morphological similarity, such as a correlation coefficient. The correlation may be treated an an edge-weight signifying the strength of a connection, but for simplicity, we will use unweighted graphs. The connectivity structure of a graph is then defined by its adjacency matrix *A* ∈ {0, 1}^*p*×*p*^, where *A*_*ij*_ = 1 if an edge exists between regions *i* and *j*, and *A*_*ij*_ = 0 otherwise. Because correlations have no directionality, *A* is symmetric.

### Estimating SCNs

#### Preprocessing and correlation

Let *N* = 1,096 denote the size of the empirical HCP dataset, and *n* the size of a (sub-)sample drawn from it. In our analysis, following (He et al., 2007; He et al., 2008), SCNs were constructed by taking structural MRI data for *n* individuals, averaging cortical thickness in each region of an atlas, and computing the sample correlations in thickness between pairs of regions.^1^ Specifically, regional cortical thickness values were assembled into a matrix *X* with *n* rows and *p* columns (brain regions, where *p* is the atlas dimensionality). Rows of *X* were Z-scored prior to computing correlations; this removes between-subject differences in global cortical thickness so that the resulting correlations reflect the spatial pattern of thickness across regions rather than overall thickness levels. We then computed Pearson *r* or Spearman *ρ* correlation for each pair of regions across individuals, yielding the sample correlation matrix 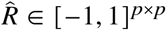.

#### Thresholding and binarization

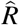 itself can be treated as an adjacency matrix, but it contains both positive and negative correlations of varying strength, and many graph-theoretic measures are defined for binary (unweighted) graphs. A common approach is to convert 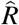 to a binary graph adjacency matrix by proportional thresholding: retaining only the strongest edges at a fixed density *δ* between zero and one (i.e., the top *δ* × 100% of edge weights). To guarantee that the resulting graph is fully connected we applied a minimum spanning tree (MST) constraint before density-based thresholding:

1. Compute the MST of the weighted graph to retain the minimal set of edges that keeps all nodes connected.
2. Add the strongest remaining edges until the target density *δ* is reached.
3. Binarize all selected edges to obtain the final estimated adjacency matrix *Â*.

We selected *δ* = 0.1 and used this SCN estimation and thresholding procedure consistently across all subsequent analyses.

### Graph theory measures

To characterise the graph-theoretic properties of the networks, we selected a number of global graph measures defined below. Clustering, average path length, modularity, and assortativity were computed using the python-igraph library (Csárdi et al., 2006); degree centrality variance was computed directly from the degree sequence returned by igraph.

We denote the degree of node *i* as *k*_*i*_ = ∑_*j,j*≠*i*_ *A*_*ij*_ (the number of connections it has with other nodes), and the shortest-path distance (minimum number of edges) between nodes *i* and *j* as *d*_*ij*_.

#### Average clustering coefficient

The clustering coefficient of a node measures the extent to which its neighbours are also connected to one another, and is therefore associated with local specialisation or segregation in brain networks. For node *i*:

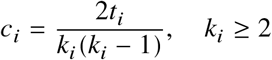

where *t*_*i*_ is the number of connections between neighbours of node *i* (i.e., triangles involving node *i*); *c*_*i*_ = 0 if *k*_*i*_ < 2. Average clustering coefficient is the mean across all nodes: 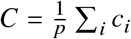.

#### Average shortest path length

Average path length captures global integration, or the efficiency of travel across the network. We define *L* as the mean shortest-path distance across all connected node pairs:

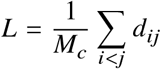

where *M*_*c*_ is the number of connected pairs.

#### Small-worldness

Small-world networks combine high local clustering with short path lengths, and is commonly studied as it indicates a balance of local specialisation and global integration in brain networks. Small-worldness is defined relative to a null model (Humphries & Gurney, 2008):

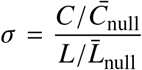

where 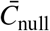 and 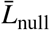 are the expected clustering coefficient and path length of an Erdős–Rényi random graph with the same number of nodes *p* and edge density *δ*. Values of *σ* > 1 indicate a small-world organisation.

#### Modularity

Modularity *Q* quantifies the degree to which the network can be partitioned into communities (modules) with dense within-module connections and sparse between-module connections. We report modularity from the Louvain community detection method (Blondel et al., 2008) with the default resolution parameter (*γ* = 1).

#### Degree variance

The degree distribution describes how connections are spread across nodes. We summarise this with the variance of the degree sequence:

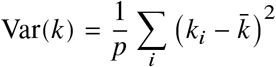

where 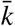 is the mean degree. High degree variance indicates the presence of hubs (nodes with disproportionately many connections) whereas low variance reflects a more homogeneous, distributed connectivity pattern.

#### Degree assortativity

Assortativity measures whether high-degree nodes tend to connect preferentially to other high-degree nodes (positive assortativity) or to low-degree nodes (negative assortativity, or disassortativity). It is the Pearson correlation of degree between the two endpoints of each edge, summed over all edges in the network:

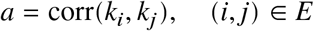

### Resampling analysis of SCN graph measures

Our empirical dataset consisted of *N* = 1,096 participants from the Human Connectome Project Young Adults preprocessed structural MRI collection. We used resampling to analyse the distribution of graph measures obtained at different sample sizes. We used two complementary resampling strategies:

1. **Bootstrapping**. Participants were resampled with replacement from the full dataset (10,000 iterations) to obtain bootstrap samples of size *N*. This yields an empirical estimate of the sampling distribution of each graph measure and provides the pool of empirical correlation matrices used as ground-truth inputs to the subsequent simulations.^2^
2. **Subsampling**. Participants were drawn without replacement in subsets of size *n* = 30 (10,000 iterations). This aims to characterise how graph measures are distributed in smaller published SCN studies.

### Simulations of SCN estimation

To understand how sample size and atlas resolution interact to affect SCN estimation quality and the sensitivity of downstream graph-theoretic comparisons, we conducted a Monte Carlo simulation (Figure 1).

**Figure 1.**
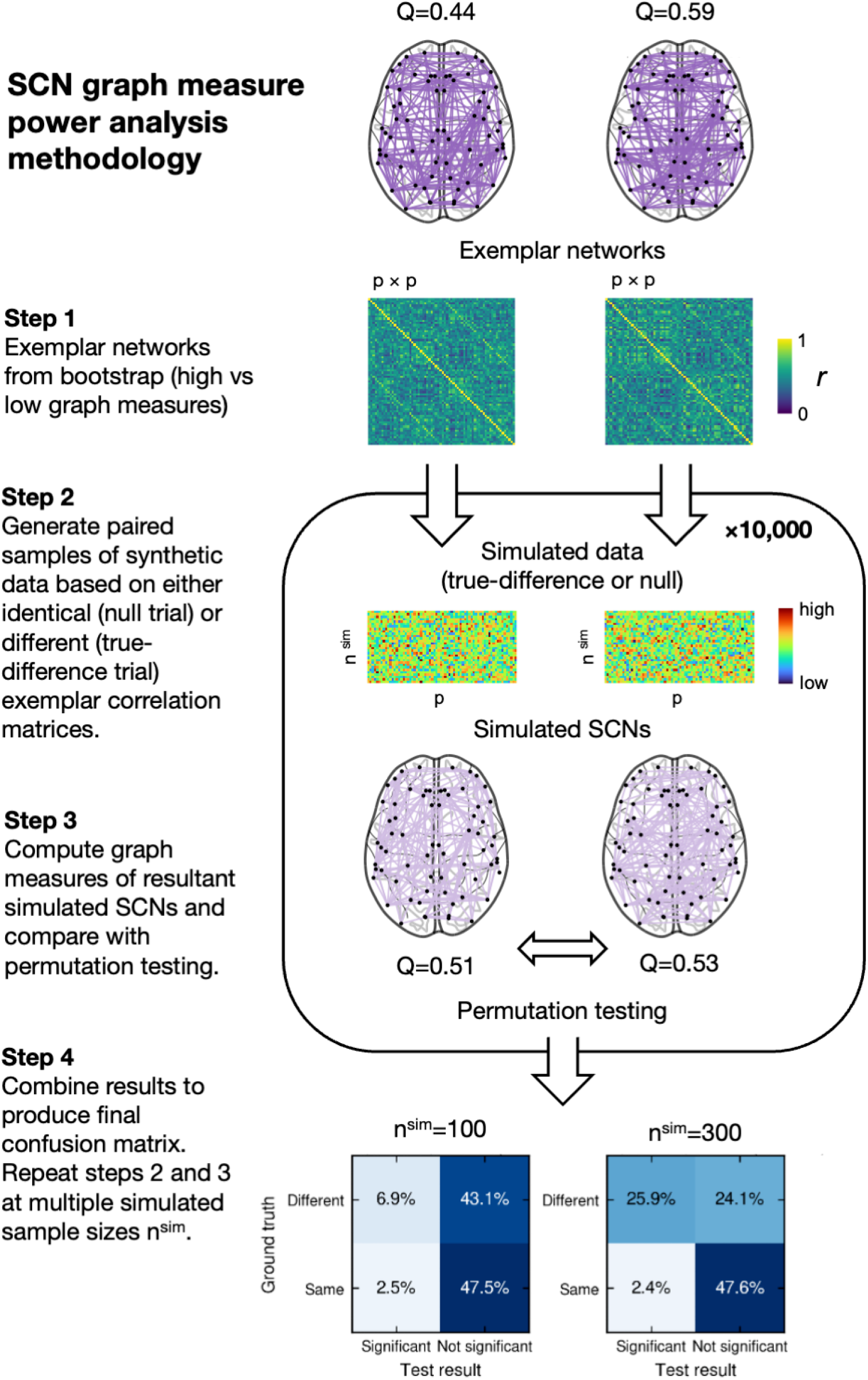
Schematic of the Monte Carlo simulation procedure. Empirical bootstrap correlation matrices serve as ground truth; multivariate normal data are generated at a given sample size, sample SCNs are estimated and thresholded, and the resulting graph measures are compared either against ground truth (reliability analysis) or between two simulated groups via permutation testing (sensitivity and specificity analysis).

#### Data-Generating Process

For each Monte Carlo trial at a given (*n*^sim^, *p*):

1. Begin with a “ground truth” correlation matrix *R*^true^ and corresponding SCN adjacency matrix *A*^true^ from the bootstrap distribution.
2. Generate *n*^sim^ i.i.d. samples *X*_*i*_ ~ *N*(0, *R*^true^), collected into an *n*^sim^ × *p* data matrix *X*.
3. Compute the sample correlation matrix 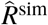 from *X*.
4. Threshold and binarise the network, as described in *Estimating SCNs* above, to obtain the simulated sample SCN *Â*^sim^.
5. Compute evaluation metrics comparing *Â*^sim^ to *A*^true^.

#### Evaluation of SCN reliability

For each combination of sample size *n*^sim^ and atlas dimensionality *p* in a predefined grid, we repeatedly generated synthetic datasets with a known “ground-truth” correlation structure and measured how accurately the sample SCN recovered it. We summarised estimation quality using two complementary measures: log-Euclidean distance and edge recall.

We use the log-Euclidean distance to quantify the distance between estimated and true correlation matrices (Arsigny et al., 2005). This metric is appropriate because correlation matrices are symmetric positive definite (SPD) and exist on a Riemannian manifold; the log-Euclidean framework respects this geometric structure. The log-Euclidean distance between 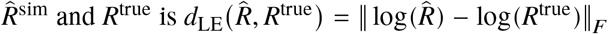.

where ‖⋅‖_*F*_ denotes the Frobenius norm. We report the Monte Carlo average of *d*_LE_ at each (*n*^sim^, *p*).

We define edge recall as the conditional probability that an edge present in the ground-truth network is included in the estimated SCN after the thresholding procedure: 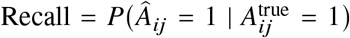. In practice, we estimate this as the average proportion of ground-truth SCN edges that are also present in the simulated sample SCN, averaged over trials for each (*n*^sim^, *p*).

We visualised each measure as a heatmap over the (*n*^sim^, *p*) grid.

### Statistical power analysis based on simulations

To evaluate the sensitivity and specificity of SCN group comparisons, we extended the simulation to mirror research practice: two groups are compared using permutation testing, and we ask how often the test reaches the correct conclusion.

#### Ground-truth correlation structures

We selected pairs of exemplar networks with corresponding empirical sample correlation matrices from the bootstrap distribution to serve as ground-truth population correlation matrices *R*^low^ and *R*^high^. For each graph measure, we defined three contrast types spanning a range of effect sizes:

- *Cohen’s d ‘large’*: a pair of networks whose metric values differed by approximately 0.8 standard deviations of the bootstrap distribution (i.e., Cohen’s *d* ≈ 0.8), selected as the nearest networks to *μ*^boot^ ± 0.8 ⋅ *s*^boot^/2.
- *Percentile*: the networks at the 1st and 99th percentiles of the bootstrap distribution.
- *Extreme*: the extreme minimum and maximum networks observed across all 10,000 bootstrap samples.

#### Permutation test simulation procedure

Group differences in SCN graph measures are commonly assessed using permutation testing, in which participant labels are exchanged between the two groups, a new SCN and graph measures computed for each permutation, and the observed difference is compared against this null distribution. To evaluate this procedure, for each combination of graph measure, contrast type, and simulated sample size *n*^sim^ ∈ {30, 50, 100, 300, 500, 1000, 3000, 5000}, we ran 10,000 independent simulation trials. In each trial we generated two groups of synthetic participants:

- *True-difference trial*: *n*^sim^ observations drawn from *N*(0, *R*^low^) and *n*^sim^ from *N*(0, *R*^high^), where *R*^low^ and *R*^high^ are the lower and upper ground-truth matrices for the given contrast.
- *Null trial*: both groups drawn from *N*(0, *R*^low^). The simulated data was then used to estimate 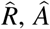 and derived graph measures for the paired groups. Each pair was compared using permutation testing (*P* = 200 label permutations), and we recorded the resulting p-value. The p-value was computed as (*c* + 1)/(*P* + 1), where *c* is the number of permutations yielding an absolute group difference at least as large as the observed difference. Pooling outcomes across all 10,000 trials, each trial was classified by its ground truth (true-difference or null) and the test decision (*p* < 0.05 or not), populating a 2 × 2 confusion matrix of true positives (TP), false positives (FP), true negatives (TN), and false negatives (FN). True-difference trials contribute the TP and FN counts (the null hypothesis is correctly rejected or missed), while null trials contribute the FP and TN counts. From these counts we estimated sensitivity (true positive rate, TPR = TP/(TP + FN)), specificity (true negative rate, TNR = TN/(TN + FP)), and the false positive rate (FPR = FP/(FP + TN) = 1 − TNR). We use the term statistical power interchangeably with sensitivity, as both are equivalent to the TPR.

#### Reporting of power and error rates

We report statistical power (the TPR at *α* = 0.05) as a function of simulated sample size for each graph measure and contrast type, since the conventional *α* = 0.05 is most representative of research practice. The TPR and FPR estimated at this same threshold together feed the false discovery rate calculation described below.

#### False discovery rate

We also report the false discovery rate, following (Ioannidis, 2005), meaning the proportion of statistically significant findings that are false positives (equivalently, one minus the positive predictive value). Because the probability that a positive test reflects a true effect depends on the prior probability that a true effect exists, which is unknown in SCN research, we report FDR as a function of prior prevalence *π*:

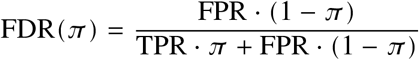

where TPR and FPR (false positive rate; proportion of null trials mistakenly identified as different) are estimated from the simulation trials. This allows readers to assess the likely FDR under their own assumptions about the prevalence of true group differences in SCN graph measures.

## Results

### Simulations of sample correlation accuracy

The first question is to what extent sample size affects the reliable estimation of SCNs. Using the full-sample correlation matrix as ground-truth, we simulated random data from a multivariate normal distribution at varying sample sizes *n*^sim^ and measured the average difference from ground-truth of the simulated sample correlation matrices and derived SCNs.

#### Log-Euclidean distance

The distance of simulated sample correlation matrices from ground truth increases with the number of regions *p*, and decreases with increasing sample size *n*, as expected *a priori* (see Vershynin, 2026, Chapter 4).

#### Edge recall

Edge recall was low at moderate to large sample sizes (Figure 3), and falls as the number of regions the cortex is parcellated into increases. Specifically, recall above 0.9 was only observed at *n* = 1000 for the most coarse atlases, the Yeo 7 and Yeo 17 resting state network parcellations (albeit with 14 and 34 nodes respectively in our analysis, due to differentiating regions across hemispheres as with the other parcellations). For atlases more commonly used in SCN analyses, such as the Desikan-Killiany cortical atlas, recall above 0.9 was only achieved at sample size of 3000, while the most high-dimensional Gordon and Glasser atlases at over 300 regions only achieved this level of SCN estimation accuracy in the *n* = 5000 simulation. On the low end of simulated sample sizes, all atlases with at least 68 regions recovered less than half of ground-truth edges at *n* = 30. Network topology is therefore unlikely to be well-preserved in SCNs estimated from small samples, especially when combined with high atlas dimensionality.

**Figure 2.**
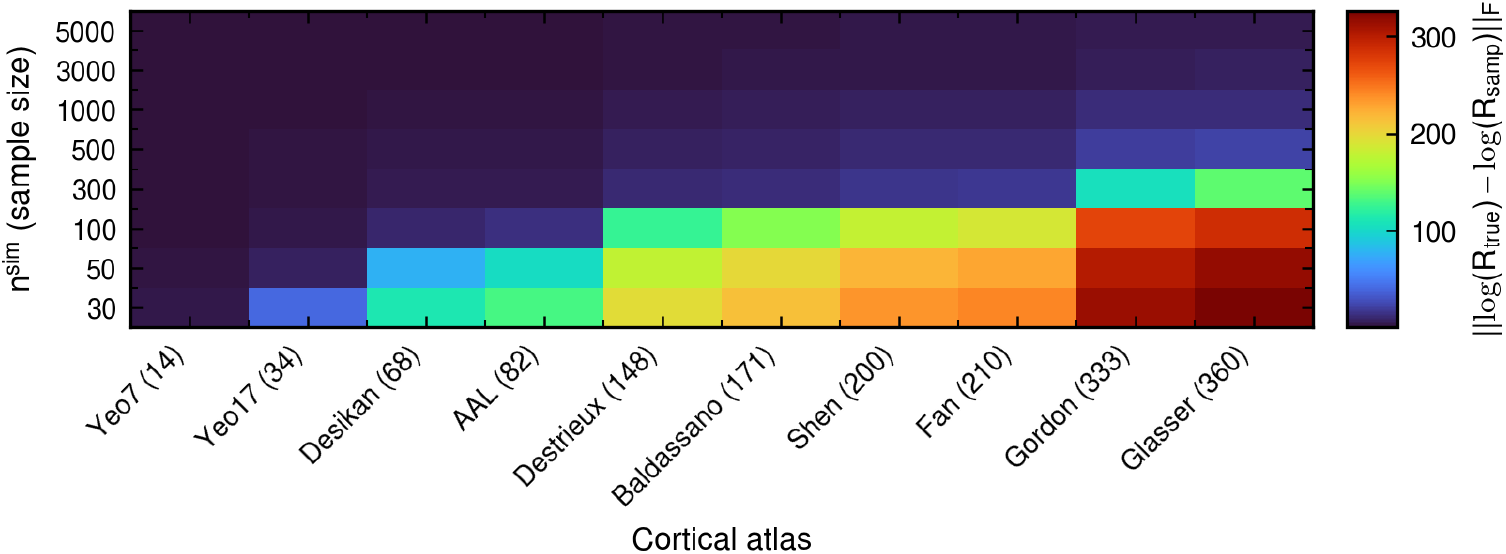
Average log-Euclidean distance of simulated sample correlation matrices from ground-truth empirical correlation matrices based on HCP data. Average *d*_LE_ of simulated sample correlation matrices from the ground-truth correlation matrix used to generate the data. x-axis: Atlas used to generate groundtruth correlation matrix (number of regions). y-axis: *n*^sim^, simulated sample size.

**Figure 3.**
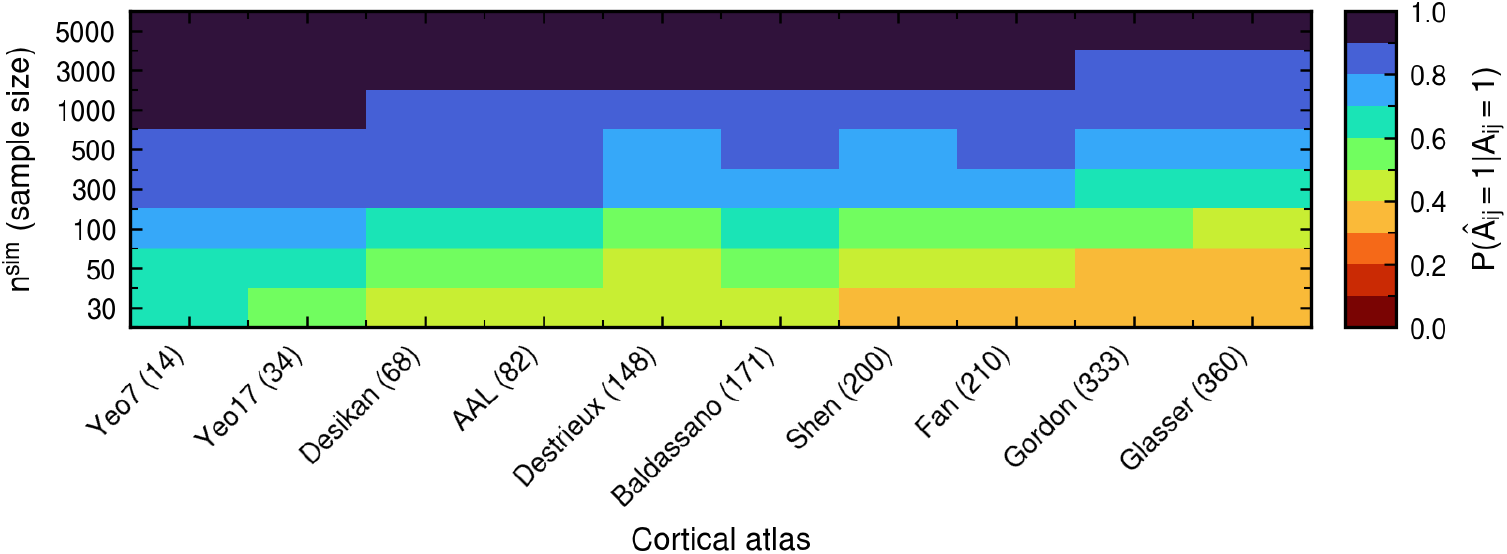
Edge recall for empirical atlas correlation matrices. Probability of an SCN edge being correctly detected as a function of sample size and atlas parcellation used. 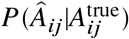 denotes the probability of an edge appearing in the simulated sample SCN, given it is in the ground-truth SCN. x-axis: Atlas used to generate ground-truth correlation matrix (number of regions). y-axis: *n*^sim^, simulated sample size.

#### Empirical analysis of SCN graph measures

Having established that sample correlation matrices diverge substantially from ground truth at small *n* and high *p*, we next examined how sample size affects the distribution of graph measures computed from empirical data. For the remaining analyses we focus on the Desikan-Killiany cortical atlas (68 regions), which is both widely used in SCN studies and, being one of the lower dimensional atlases in our study, a relatively favourable case for reliable estimation; the results that follow should therefore be regarded as a best case relative to the finer parcellations also in common use.

#### Bootstrap sample distributions

Bootstrapping the HCP Young Adults preprocessed structural MRI data (Figure 4), we observe generally ‘small-world’ cortical thickness SCNs, according to the (Humphries & Gurney, 2008) definition (*σ* > 1). Crucially, we observe that the subsampling (n=30) distribution of graph measures differs substantially from the bootstrap distribution, with all the graph measures we investigated being biased downwards in smaller samples (although reduced path length would translate to increased ‘network efficiency’ in small *n* networks, for instance). The distribution of graph measures for the small samples also exhibits increased variance.

**Figure 4.**
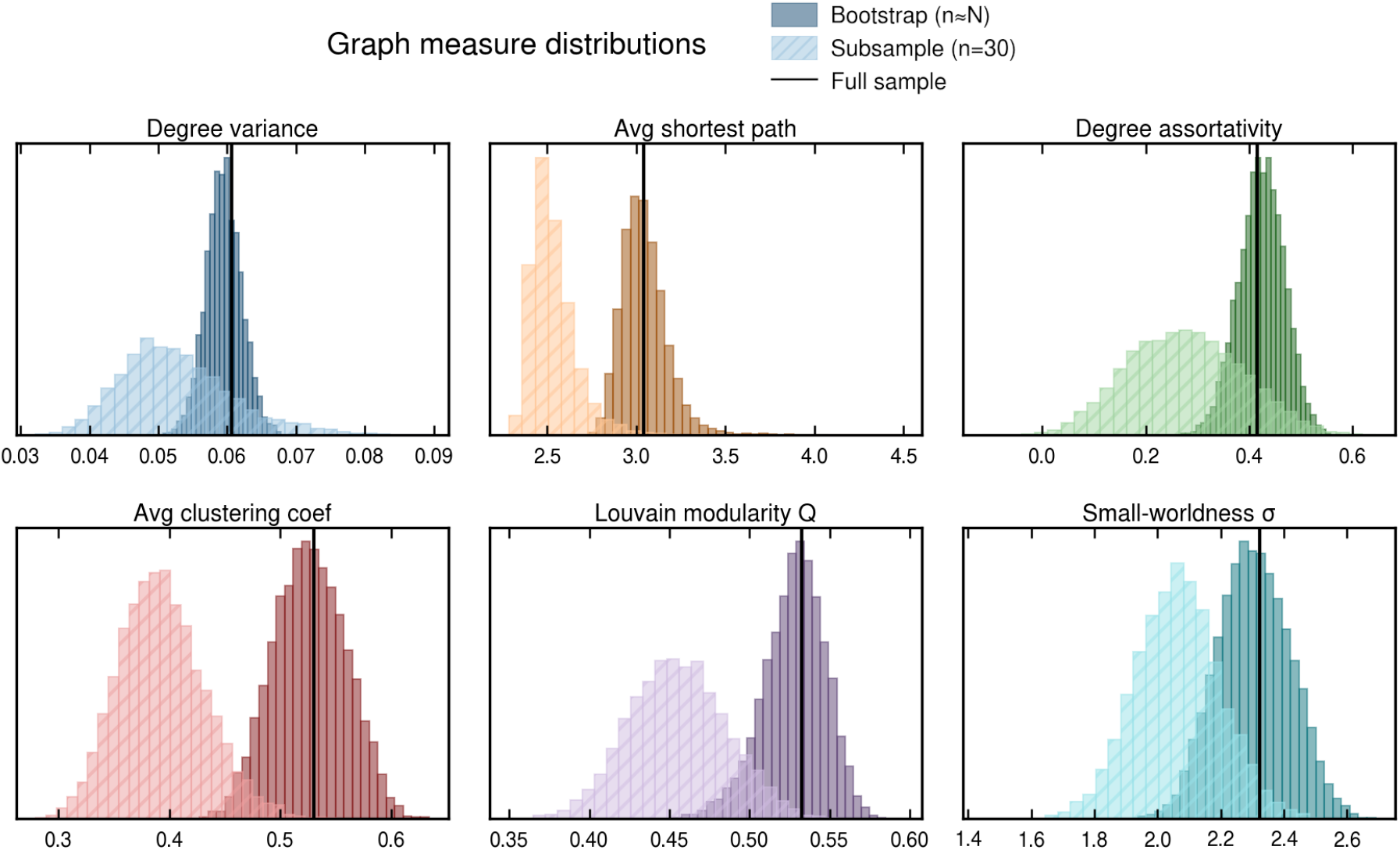
Distributions of 6 graph measures from bootstrapping empirical data at matched simulated sample sizes (large: *n* ≈ 1096; small: *n* = 30) using the Desikan-Killiany atlas (*p* = 68). The darker histogram shows the distribution of graph measures for the large-sample SCNs, while the transparent, hatched histogram shows the distribution for the small-sample SCNs. The solid black line indicates the network obtained from the original full sample.

### Sample size relationship with graph measures

Observing that the distribution of graph measures differs greatly between small and large samples, we next conducted bootstrapping across a range of *n* to observe the precise relationship between sample size and graph measures. For each *n* at 25 log-spaced intervals between 10 and 1000 we drew 10,000 samples with replacement, and plotted the median and interquartile range of resulting SCN graph measures in Figure 5. Interestingly, for most graph measures the relationship is nonlinear, reaching a minimum for sample sizes around 20 or 30, before increasing until *n* ≈ 1000. Notably, the difference in graph measure distributions between e.g., *n* = 30 SCNs and *n* = 100 SCNs is substantial and therefore even this relatively small difference in sample sizes between groups will lead to severely biased comparisons.

**Figure 5.**
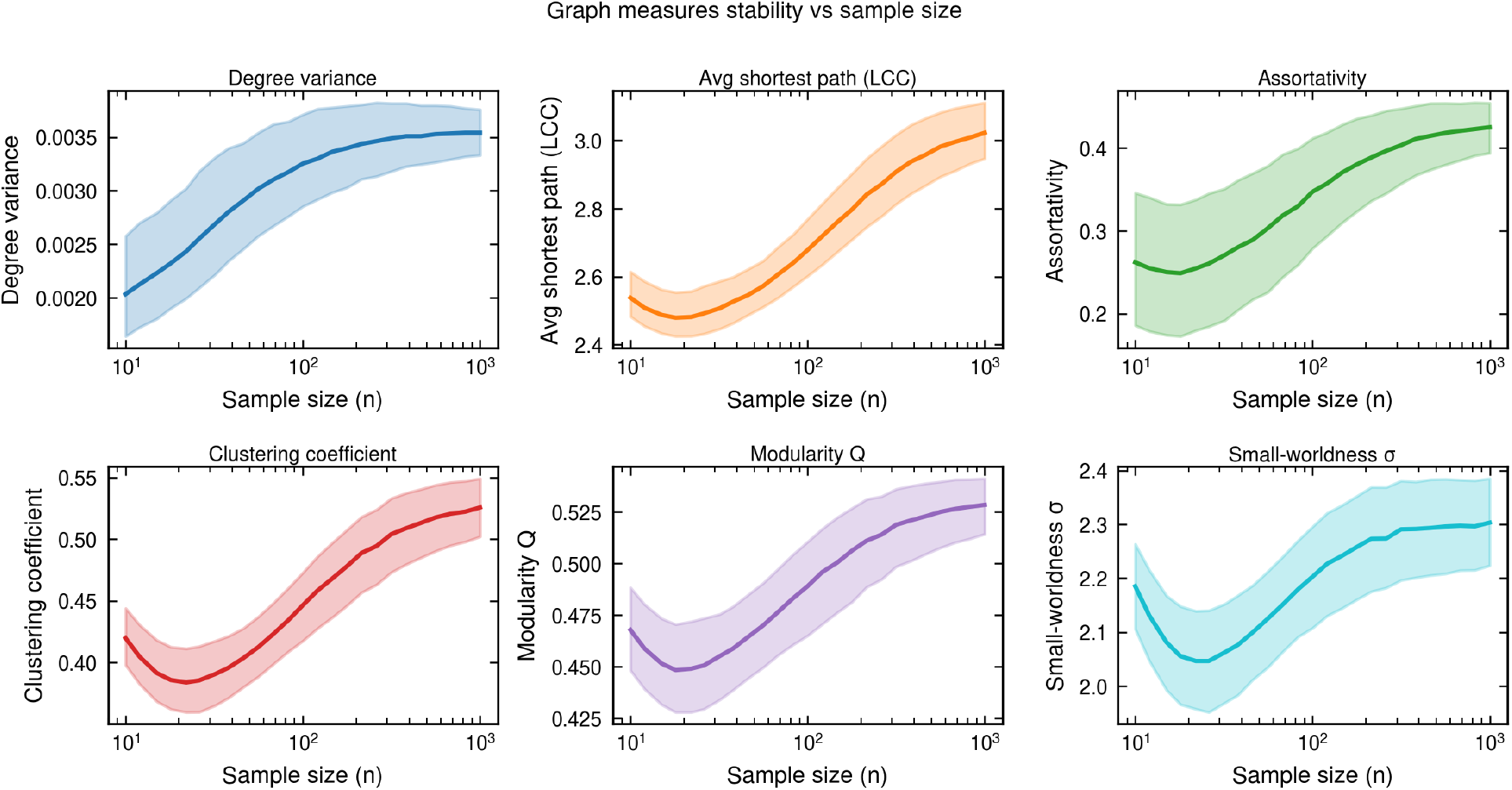
Results of bootstrapping and calculating graph measures across a range of *n* using the Desikan-Killiany atlas (*p* = 68). Line: bootstrap median. Shaded area: interquartile range.

### Statistical power analysis based on simulations

The preceding analyses demonstrate that SCNs are unreliably estimated at common sample sizes/atlas dimensionalities, and furthermore that small samples yield biased graph measures. We next asked whether standard permutation testing can reliably detect true differences in groups when sample sizes are matched, or whether the noise revealed by our earlier simulations in small-sample SCN edges leads to low statistical power and unacceptable false discovery rates at common sample sizes.

The permutation testing simulations (Figure 6) demonstrate that the statistical power achieved for a conventionally ‘large’ effect size (*d* = 0.8), defined according to the variance observed in the bootstrap distribution (Figure 4), is well below 0.8 at all simulated sample sizes tested. The best performing graph measure for statistical power was by far the average clustering coefficient, which reached a power of approximately 0.6 at *n* = 5000. Power above 0.8 was achieved only for much larger effect sizes, using either the 1st and 99th percentile of the bootstrap distributions (‘percentile’ in Figure 6) or the actual minimum and maximum of each graph measure observed in the 10,000 bootstrapped SCNs (‘extreme’ in Figure 6), equivalent to very large Cohen’s *d* of 4+ and 6+ respectively. Nonetheless, even in the case of average clustering coefficient, a sample size of 300+ was required to achieve power of 0.8 in the ‘extreme’ effect size condition. Therefore for permutation testing of SCN graph measures to reach conventionally accepted levels of statistical power, even at effect sizes far beyond normal, per-group sample sizes must be at least in the hundreds or thousands.

**Figure 6.**
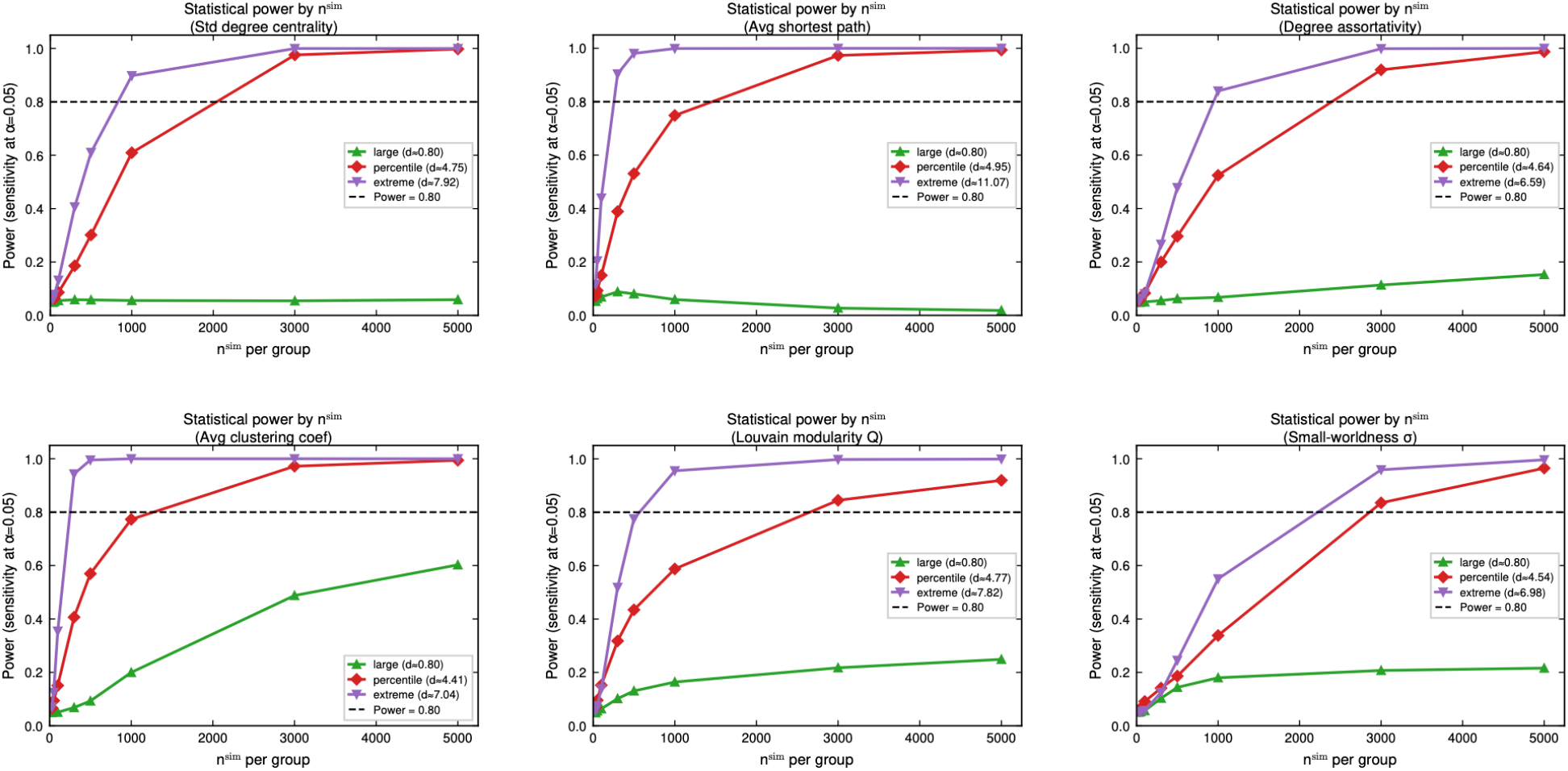
Statistical power achieved across simulated sample sizes for each of the six graph measures using the Desikan-Killiany atlas (*p* = 68). Line style indicates the ground-truth difference used for the simulation. Green line w/ triangle marker: Cohen’s *d* ≈ 0.8. Red line w/ diamond marker: 1st and 99th percentile of bootstrapped graph measures. Purple line w/ inverted triangle marker: Minimum and maximum of bootstrap graph measure distribution.

### Estimates of false discovery rate according to prior probability of an effect

Given the estimates of statistical power/sensitivity obtained in the permutation testing simulations, we infer the false discovery rate (FDR) that will result across the range of true-effect rates *π*, i.e., the prior probability of a true effect actually existing (following (Ioannidis, 2005)). Using the FDR expression defined in the Methods, this is the proportion of significant tests that are false positives.

Figure 7 shows the expected FDR across sample sizes and true-effect rates *π*. At *n*^*sim*^ = 30 across all three effect sizes, the false discovery rate is almost perfectly proportional to *π*, indicating that permutation testing for differences in modularity was no better than chance at this sample size. When *π* is low at the Cohen’s *d* ‘large’ effect size, such as at *π* = 0.15 or 15% of group comparisons tested having a ground-truth ‘large’ difference in modularity, the majority of significant tests would be false discoveries even at *n* = 5000. While the true level of *π* is in practice never known, it should be assumed that the larger the effect size, the lower the rate of ground-truth differences of that magnitude will be. Therefore, quite conservatively assuming *π* < 0.25, we can predict that the majority of ‘significant’ graph measure differences in SCNs are false positives when *n* ≤ 100.

**Figure 7.**
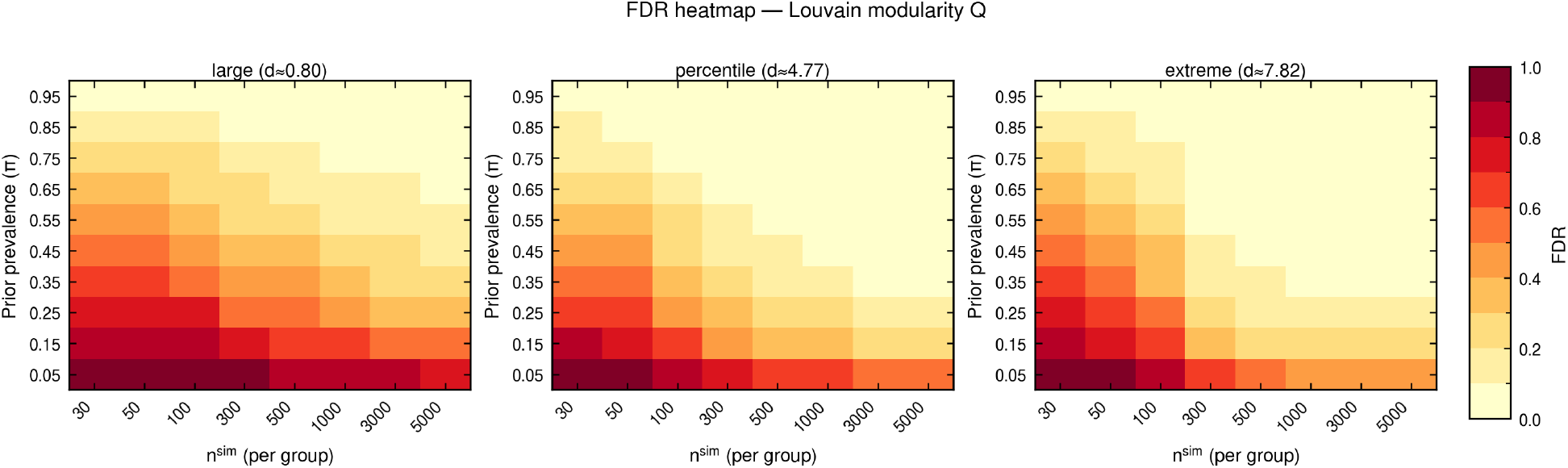
Expected false discovery rate (FDR) for three simulated SCN effect sizes of modularity Q. The prior prevalence of true effects *π* is allowed to vary between 0.05 and 0.95 of statistical tests, while *n*^*sim*^ varies between 30 and 5000. The heatmap colour represents the expected FDR, given the sensitivity and specificity at each *n*^*sim*^ measured in the permutation testing simulation. y-axis: True-effect rate *π*. x-axis: *n*^sim^, simulated sample size.

## Discussion

We set out to characterise the statistical validity of graph-theoretic comparisons of structural covariance networks, estimating the reliability of SCN edges, the dependence of graph measures on sample size, and the statistical power and false discovery rate of permutation testing across sample sizes and atlas dimensionalities. We find that SCNs are unreliably estimated at the sample sizes and parcellation resolutions common in the literature, that small samples systematically bias graph measures, and that permutation testing is no better than chance at *n* = 30 — such that, under reasonable assumptions about the prior prevalence of true effects, the majority of significant graph-measure differences at *n* ≤ 100 are likely to be false positives.

### Reliability of SCN estimation

Our simulation results demonstrate that the reliability of SCN edges is especially low for high-dimensional atlases, requiring correspondingly larger sample sizes for robust estimation. Studies with smaller sample sizes should therefore restrict analyses to networks with fewer nodes. The relationship between parcellation granularity and reliability is shown clearly in Figure 3: edge recall falls below 50% for SCNs estimated on commonly used atlases (68+ regions) at sample sizes often seen in published studies to date (*n* < 50).

Importantly, measurement error does not appear to be the primary driver of unreliability. In analyses not reported here, adding simulated measurement error scaled to test-retest estimates led to only slight worsening of performance compared to the dominant effect of sampling variance (not shown). This suggests that the fundamental constraint is the ratio of observations to parameters being estimated (the ‘curse of dimensionality’, (Altman & Krzywinski, 2018)). As a rule of thumb, we propose considering the number of free parameters in a correlation matrix of *p* variables, equal to 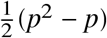. For the Glasser atlas with 360 bilateral nodes, this amounts to 64,620 individual correlations to be estimated – far exceeding typical sample sizes. Alternatively, consider that according to theoretical results the Frobenius norm error of sample covariance matrices for normally distributed data scales as 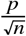, suggesting that the required sample size for reliable SCN estimation scales quadratically with respect to the atlas dimensionality/number of brain regions (Vershynin, 2026, Chapter 4). However, this scaling law only holds for Gaussian or sub-Gaussian distributed data, meaning that real data with heavy tails and outliers will likely scale worse in practice. This consideration also applies to our simulations, which use a multivariate normal distribution to generate data and are therefore more forgiving than real data is likely to be.

In this study, we treated parcellations as bihemispheric, reflecting common practice. In exploratory analyses not reported here, we found that without normalising cortical thickness data across both hemispheres, some SCNs were highly biased to one hemisphere (not shown). Averaging parcellations across hemispheres would both eliminate this effect and reduce dimensionality, which should improve SCN reliability given the dependence of reliability on dimensionality shown in Figures 3 and 2.

### Dependence of graph measures on SCN sample size

As shown in Figure 4 and Figure 5, our bootstrapping and subsampling analysis indicates that the distribution of graph measures is highly sensitive to the sample size. Specifically, random edges introduced by sampling noise will tend to reduce measures of segregation such as clustering coefficient and modularity, while inflating measures of integration such as global efficiency. Therefore, studies that compare groups of different sizes (such as when a small group of patients is compared to a large group of healthy controls) are likely to find significant effects simply as a result of this disparity – the group with smaller *n* will appear to have a more integrated, less modular network topology regardless of any true biological difference. Finally, as the variance of graph measures varies in some cases with sample size, the exchangeability assumption for permutation testing may be violated, leading to an inflated FPR (Huang et al., 2006).

### Statistical power and false discovery rate of SCN graph-theoretic comparisons

As shown in Figure 6, the statistical power of graph measure comparisons was well below 0.8 for all sample sizes and graph measures tested, when based on a ground-truth difference of *d* = 0.8. High statistical power was achievable for all graph measures when the ground-truth networks were the 1st and 99th percentile of the bootstrap graph measure distribution (‘percentile’ in Figure 6) only when *n* ≥ 3000. When the ground-truth difference was the minimum and maximum of the entire 10,000 sample bootstrap distribution (‘extreme’ in Figure 6), a simulated sample size between 300 and 3000 sufficed to reach a power of 0.8, depending on the specific graph measure. While average shortest path length and average clustering coefficient exhibited higher power than other graph measures, their normalized ratio *σ* required the largest sample size to reach a power of 0.8 out of all graph measures tested, suggesting that the combination of integration and segregation embodied in ‘small-world’ networks is more difficult to reliably detect.

Converting the statistical power estimates at each *n*^sim^ to FDR, contingent on the prior probability of a true effect *π*, reveals that how meaningful ‘significant’ differences in SCN graph measures are depends not only on the test’s sensitivity, but also our prior expectations about the likelihood of these differences. Given the magnitude of the ‘percentile’ and ‘extreme’ effect sizes (a 1-in-50 difference and a 1-in-10,000 difference respectively), it is reasonable to assume that they occur infrequently – but we must also account for researchers’ choice of topic. If researchers choose topics such that there is a 50/50 chance of an ‘extreme’, 1-in-10,000 difference in a single graph measure (i.e., not considering the problem of multiple comparisons), a sample size of 500–1000 may suffice to reduce the chance of a false discovery below 10%, provided one can match sample sizes between the groups. However, if there is no particular reason to expect differences in graph-theoretic properties of SCNs (e.g., *π* = 0.05), false discoveries may compose the majority of significant findings even with a per-group sample size of 5000+. For this reason, it is especially important to carefully consider our reasons for hypothesising graph-theoretic differences ((Bullmore & Sporns, 2009)), which usually relate to the information processing properties of brain networks. SCNs reflect ‘mutually trophic’ associations between brain structure in different regions (Mechelli et al., 2005), which is less easily related to information processing than measures of connectivity from functional imaging.

Our statistical power analysis was confined to the Desikan-Killiany atlas, as this is commonly used in SCN studies. Based on the earlier simulation across multiple atlases shown in Figures 2 and 3, statistical power will only be worse for atlases with more regions than Desikan-Killiany. As suggested above, estimating SCNs with fewer nodes is advisable to avoid the curse of dimensionality – but this does not preclude the use of atlases that define many regions, such as Glasser. Rather than including every region in a brain-wide atlas, researchers may instead select a subset of regions of interest to compute an SCN, as in the early work of (Mechelli et al., 2005).

### Limitations

We derived Cohen’s *d* estimates by assuming that compared groups’ standard deviation in graph measures is equal to that of the healthy control population in the HCP sample. This is the standard assumption underlying the application of Cohen’s *d*, and in cases where standard deviation differs between groups, the alternative measure Glass’ Δ uses the control group standard deviation. Therefore, although this method of estimating the ‘large’ effect size leads to quite small nominal differences in graph measures, it should be a reasonable proxy for real-world effect sizes. However, it should be considered that in some conditions, for instance especially neurodegenerative diseases that cause severe coordinated brain atrophy, extremely large effect sizes *d* ≫ 2 may be expected. For this reason, we include comparisons of the ‘percentile’ (4 < *d* < 5) and ‘extreme’ (6 < *d* < 11) effect sizes, but we note that even these profoundly divergent cases, the required sample size is still large relative to most published SCN studies.

Our simulation is less challenging than real-world SCN estimation. This is because our simulated data is normally distributed, while real-world cortical thickness and morphometry data tends to deviate from normality, exhibiting heavy tails, skewness, and outliers that make covariance estimation more challenging. In addition, correlated measurement errors due, e.g., to head motion that introduce spurious dependencies between regional cortical thickness values may vary between groups of interest and thus increase the rate of false positives (Read-Tannock et al., 2026). Therefore, the estimates of sensitivity we obtained should be viewed as an upper bound on the true sensitivity of graph-theoretic SCN comparisons, and thus the false discovery rate may in fact be higher than our results indicate.

We have also not considered node-wise graph measures such as betweenness centrality used to identify network hubs in many studies, but these will of course be even more sensitive to correlation matrix misestimation and consequent false positive edges.

### Implications and future directions

False positives are rare using the permutation testing approach (in our simulation as expected, FPR ≈ *α*), but in the presence of high noise in covariance matrix estimation, true positives are rare also. In other words, the specificity of permutation testing is high, but the sensitivity is low. Consequently, though there may be a true difference in graph measures between groups, a prohibitively high sample size is needed in order to reliably detect those differences at commonly used atlas dimensionalities. The sample size requirement is much larger than for simple statistical tests (e.g., a t-test of group means). This is because there is a high-dimensional estimator (the correlation matrix) in between the sampled data and the test statistic. Large open datasets such as the UK Biobank are consequently the best test-beds for high-dimensional SCN research, and could be productively employed to obtain better estimates of population variability and for replication studies (Kharabian Masouleh et al., 2019; Madan, 2022).

The practice of reporting the number of permutations used for the hypothesis test is worthwhile, but very high numbers of permutations only increase the precision of the p-value obtained, and cannot compensate for excessive noise in the test statistic. In the case of graph measures derived from SCNs, the sample size *n* and atlas dimensionality *p* are the crucial factors determining how stable the covariance/correlation matrix will be, and thus how meaningful derived graph measures are and how effective permutation testing will be.

One method for reducing the influence of false positive or random edges present in small sample SCNs is to evaluate graph measures across a range of proportional thresholds, and then comparing AUC or identifying significant effects in clusters of thresholds (Drakesmith et al., 2015). This should increase the robustness of graph-theoretic comparisons, but we did not assess to what degree as it would increase the time required for simulations multiplicatively.

A natural extension of this work is to apply similar validation to the growing family of alternative structural co-variance methods. As noted in the Introduction, numerous variants have been proposed, including individual-level SCNs based on regional morphometric similarity (Kong et al., 2015; Tijms et al., 2012; Wee et al., 2012), individual differential SCN (IDSCN; (Liu et al., 2021)), morphometric similarity networks (MSN; (Seidlitz et al., 2018)), morphometric inverse divergence networks (MIND; (Sebenius et al., 2023)), and causal SCNs using Granger causality on observations ordered by a variable of interest (Zhang et al., 2017). Some, such as MSN and MIND, leverage multivariate morphometric data and may benefit from improved power relative to univariate approaches. Systematic benchmarking of reliability and false discovery rates across this family of methods would be valuable as they see increased application in research.

Additionally, generative models for structural covariance networks could provide informative prior expectations that may serve as improved null distributions (Betzel & Bassett, 2017). Shrinkage regularisation of the sample correlation matrix is another promising avenue for improving statistical power (Brier et al., 2015). (Honnorat & Habes, 2022) investigated shrinkage toward the identity matrix, but because the identity matrix in linear combination with the sample correlation affects all correlations equally, this does not alter the rank ordering of correlations and thus only benefits absolute thresholding approaches. A more informative shrinkage target may be the cortical distance matrix (Euclidean or geodesic), reflecting the robust finding that structural covariance is stronger at shorter distances (Alexander-Bloch et al., 2013).

Further work is also needed to understand how edge-level errors propagate to graph measures. It is possible that low edge recall severely degrades the reliability of graph measures if false positive edges are randomly distributed – however, false positive edges included in the SCN are likely similarly distributed to true edges, potentially limiting the problem especially when analysed over a range of thresholds. Finally, while our simulations suggest that measurement error scaled to test-retest estimates (Iscan et al., 2015) contributes less to SCN unreliability than sampling variance, a systematic treatment of head motion effects on SCN estimation would complement existing work (Pardoe & Martin, 2022).

## Conclusions

In summary, our analysis indicates that graph-theoretic comparisons of structural covariance networks are sensitive to sample size and parcellation resolution, with false discovery rates elevated at sample sizes common in the literature. These findings highlight the value of larger samples, use of coarser or partial anatomical atlases, and explicit reliability assessment when interpreting group differences in SCN topology. Under reasonable assumptions about the prior prevalence of true effects, a substantial proportion of published findings at small sample sizes are likely to be false positives; we hope this work encourages continued methodological development and helps researchers design SCN studies with appropriate statistical power.

## Footnotes

1 SCNs of this kind are, like functional connectivity networks, based on correlation matrices. Networks estimated by thresholding correlation matrices (correlation graphs) have specific properties, including elevated transitivity, clustering coefficient and smallworldness relative to random graphs (Hlinka et al., 2017; Zalesky et al., 2012).

2 Conditional on network methodological choices such as threshold, choice of correlation measure, etc.

